# *barcodetrackR*: an R package for the interrogation of clonal tracking data

**DOI:** 10.1101/2020.07.23.212787

**Authors:** Diego A. Espinoza, Ryland D. Mortlock, Samson J. Koelle, Chuanfeng Wu, Cynthia E. Dunbar

**Author notes:** Equal contributions (Diego A. Espinoza, Ryland D. Mortlock). Corresponding author, Correspondence should be addressed to: Cynthia E. Dunbar, Translational Stem Cell Biology Branch, NHLBI, NIH, Building 10 CRC, Room 5E-3332, 10 Center Drive, Bethesda, Maryland 20892.

## Abstract

Clonal tracking methods provide quantitative insights into the cellular output of genetically labelled progenitor cells across time and cellular compartments. In the context of gene and cell therapies, clonal tracking methods have enabled the tracking of progenitor cell output both in humans receiving cellular therapies and in corresponding animal models, providing valuable insight into lineage reconstitution, clonal dynamics, and vector genotoxicity. However, the absence of a toolbox by which to interrogate these data has precluded the development of standardized analytical frameworks within the field. Thus, we developed *barcodetrackR*, an R package that provides users with tools for the analysis and visualization of clonal dynamics across time and cellular compartments in clonal tracking experiments. Here, we demonstrate the utility of *barcodetrackR* in exploring longitudinal clonal patterns and lineage relationships in the context of a number of clonal tracking studies of hematopoietic stem and progenitor cells (HSPCs) in humans receiving HSPC gene therapy and in animals receiving lentivirally transduced HSPC transplants.

## INTRODUCTION

Genetic labelling permits quantitative tracking of clonal progeny via high-throughput sequencing (clonal tracking) and provides opportunities to interrogate clonal dynamics in a number of *in vitro* and *in vivo* contexts. The two most common clonal tracking approaches, cellular barcoding and viral integration site recovery, have been primarily leveraged to interrogate hematopoietic stem and progenitor cell (HSPC) or immune cell dynamics both in model animals(1–5) and in humans(6,7). In these methodologies, integrating retro- or lentiviruses are used to transduce individual HSPCs or other target populations such that individual cells each contain a unique, permanent genetic tag or integration site label that can be recovered from progeny cells via high throughput sequencing (**Fig. 1**). Measurement of each label’s abundance in the pool of all recovered labels is directly associated with the abundance of that clone within the labelled population being assayed, for instance T cells, B cells or myeloid cells. These lineage abundance measurements can provide insights not only into the bias, stability, and ontogenetic relationships of HSPCs(8), but also into the dynamics of clones within cell populations whose abundances are largely independent of HSPC behavior, such as certain T cell(9) and natural killer (NK) cell subsets(10). Furthermore, such clonal tracking methods have also been leveraged to provide valuable insight into the clonal dynamics of cancer progression(11), *in vitro* differentiation(12), and clonal dynamics of CAR-T cells(13).

**Figure 1:**
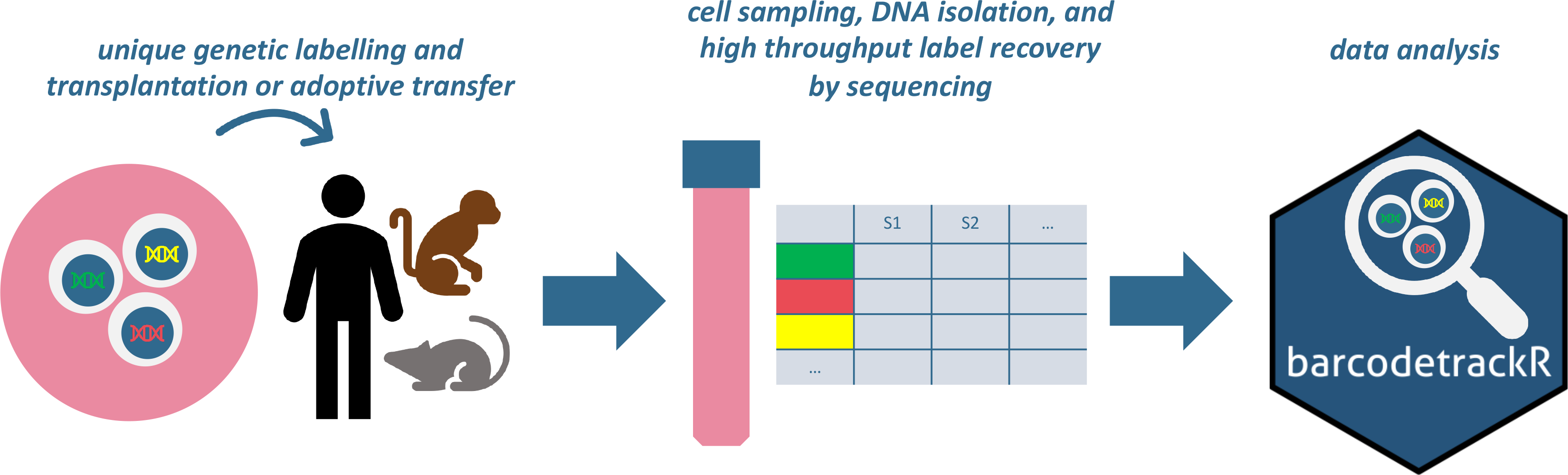
Clonal tracking experimental design. Clonal tracking experiments generally follow the depicted framework, which encompasses, in order, the genetic labelling of cells to create an integrated DNA tag, transplantation or adoptive transfer of these cells into a recipient, acquisition of cellular progeny from the recipient following transplantation or adoptive transfer, genetic tag retrieval from progeny cells followed by high-throughput sequencing, algorithmic quantification of detected individual unique tags, and finally, downstream analyses, where the *barcodetrackR* toolkit can be utilized.

Given the diversity of labelling and recovery strategies, as well as underlying differences in vector and barcode constructs, a number of approaches for recovery of sequences from raw sequencing data and determination of “true” integration site or genetic tag from sequencing artifacts or other confounders have been developed and are largely approach-dependent(14–17). However, tools with which to perform downstream analyses of the clonal abundances determined by these pipelines have not been published or made publicly available; as a result, flexible open-source tools, such as those that exist for single-cell RNA-sequencing(18,19) have been sought after by those in the clonal tracking field in order to derive biological meaning in an accessible manner from these large datasets(20). Such tools would also allow direct comparisons across datasets or meta-analyses.

Here, we present our open-source R package, *barcodetrackR. barcodetrackR* encompasses a variety of flexible tools that can provide insights into clonal dynamics and the relationships between cellular compartments starting with clonal abundance data. We illustrate the utility of *barcodetrackR* by analyzing publicly available clonal tracking datasets from studies in lentivirally transduced non-human primates(8,10,21), xenograft mice transplanted with human cord blood cells(22) and blast cells(23), and lentiviral gene therapy patients(6,24). More details on each dataset and access paths are summarized in Table 1.

**TABLE 1:**
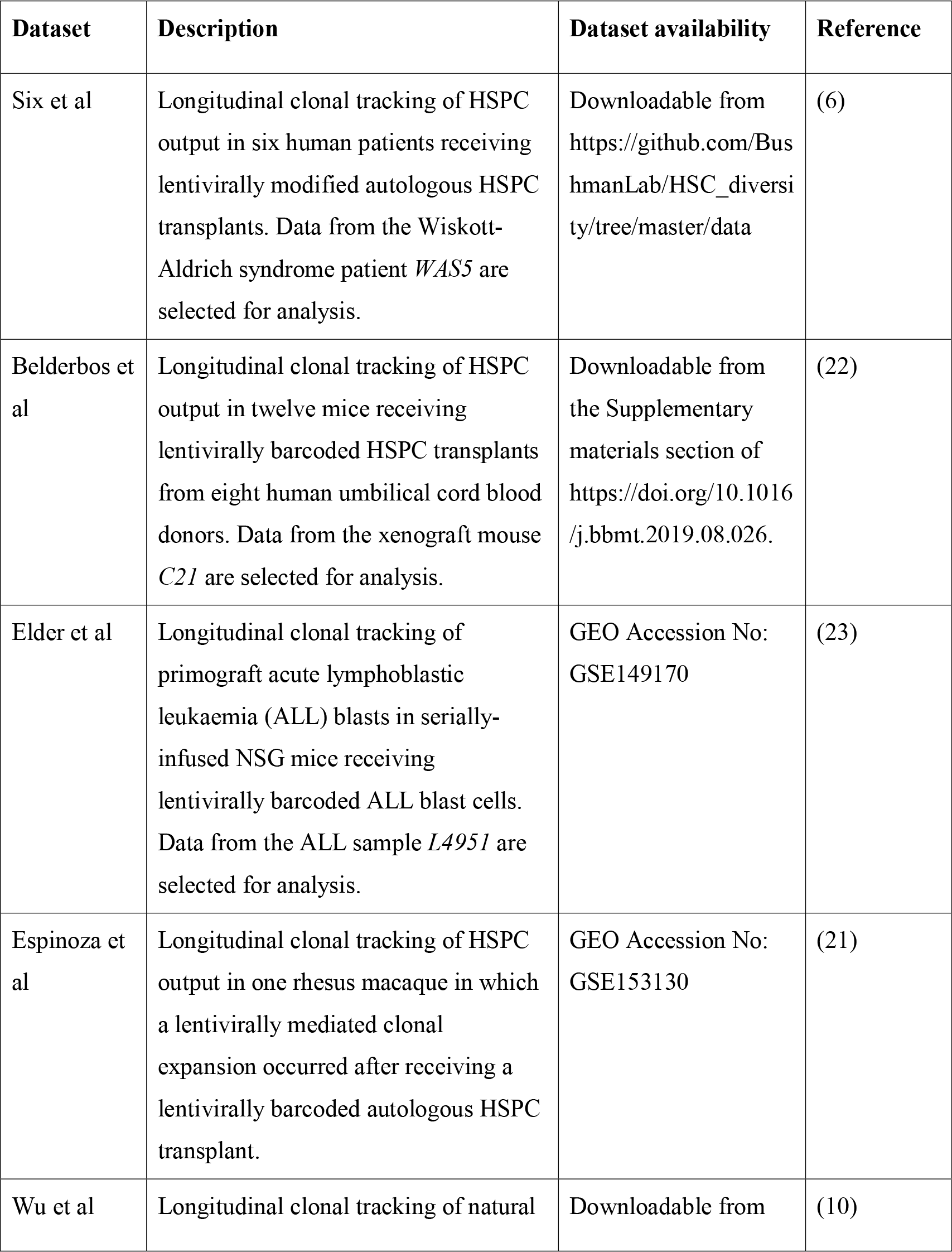
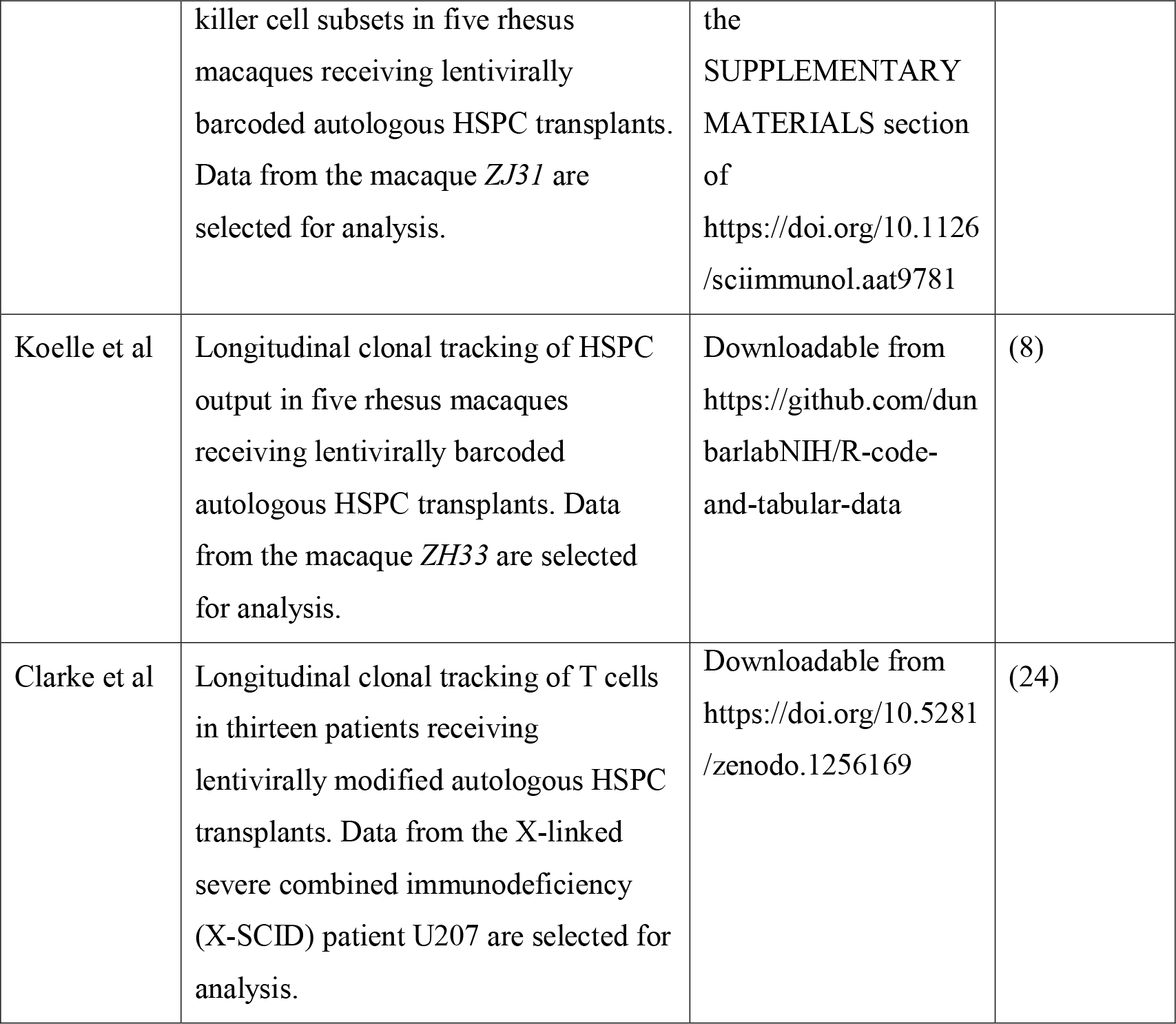
EXAMPLE DATASETS

## RESULTS

### Inferring lineage relationships based on global clonal distributions

Pairwise comparisons of clonal abundance profiles in clonal tracking data provide insight into the relationships between upstream progenitor pools across cellular compartments. Here, we use *barcodetrackR* to determine and visualize the correlation values and dissimilarity indices (**Fig. 2**) between samples from three clonal tracking datasets (Six et al, Belderbos et al, Elder et al, Table 1) as a means to interrogate the similarities of upstream progenitor pools contributing across cellular compartments. The Six dataset contains individual viral integration site read counts from longitudinally collected patient T cell, B cell, granulocyte (Gr), monocyte (Mo), and natural killer cell (NK) samples following autologous lentiviral HSPC gene therapy. We find that the Gr and Mo samples share high correlation with one another, while the T cell, B cell, and NK samples show lower correlation with samples from other lineages, but high correlation between different timepoints within the same lineage (**Fig. 2A**). A similar pattern is observed when plotting the Bray-Curtis dissimilarity indices between samples from the Six dataset, projected into two dimensions using principal coordinates analysis (PCoA), where the first axis of variation separates NK cells from other lineages based on their clonal abundances, and the second separates T cells, B cells, and myeloid (Gr and Mo) cells (**Fig. 2B**). These analyses demonstrate that the myeloid lineages are closely coupled and thus likely arise from shared pathways originating from the same HSPC pool, in comparison to disparate generation of mature T, B, and NK lineages.

**Figure 2:**
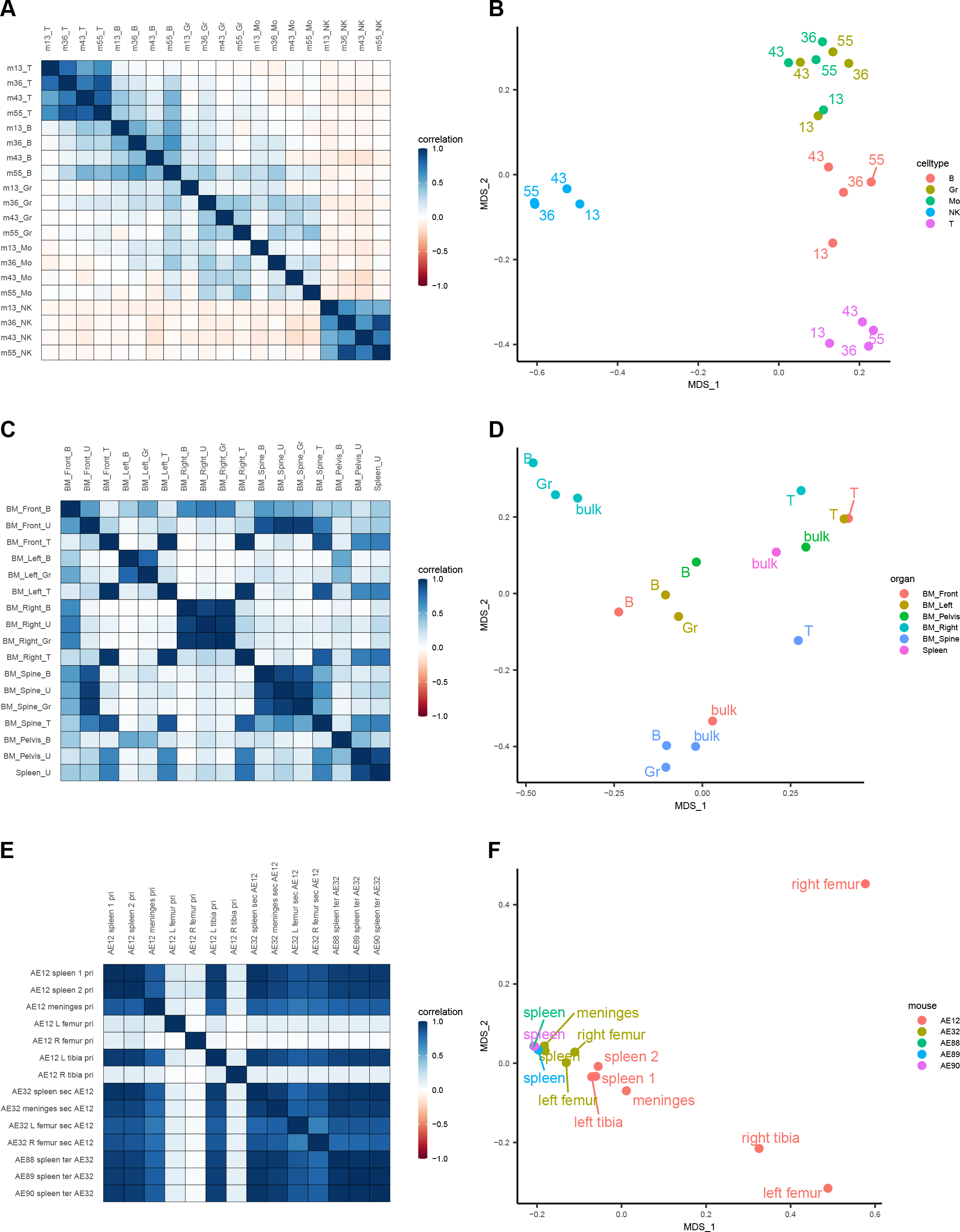
Global clonal distributions. Global clonal distributions in the Six, Belderbos, and Elder datasets. Pairwise Pearson correlation plots between longitudinal samples from the Six dataset (**A**). Row and column labels indicate months post-transplant (*m*) and cell type (*Gr, Granulocyte; Mo, Monocyte; NK, Natural Killer cell*). Bray-Curtis dissimilarity indices between samples from the Six dataset (**B**), grouped by cell type and labeled based on months post-transplant. The x and y-axis represent the two main axes of variation after conducting principal coordinate analysis on the Bray-Curtis measures of dissimilarity (*MDS*). Pairwise Pearson correlation plots between samples from different anatomical sites of a single transplanted mouse at euthanasia from the Belderbos dataset (**C**). Row and columns labels describe the anatomical site (*BM, Bone Marrow*) followed by the cell type (*U, Unsorted samples*). Bray-Curtis dissimilarity indices between samples from the Belderbos dataset (**D**) grouped by the anatomical site and labeled by cell type. Pairwise Pearson correlation values between samples of the same set of serial xenograft transplants from the Elder dataset (**E**). Row and column labels describe the animal code (*e.g. AE12*), followed by the anatomical site, then the serial transplant designation (*pri, Primary; sec, Secondary, ter, Tertiary*), followed by the donor animal code if it is a *sec* or *ter* sample. *AE12* is the primary recipient of ALL blast cells, *AE32* is the secondary recipient receiving cells from the primary animal *AE12*, and *AE88, AE89, AE90* are tertiary recipients receiving cells from *AE32*. Bray-Curtis dissimilarity indices between samples from the Elder dataset (**F**) colored by the mouse of origin and labeled by the anatomical site.

These pairwise measures can be also used to compare clonal abundances across anatomical compartments. The Belderbos dataset contains clonal abundance information from a number of sorted and unsorted immune cell samples from bone marrow (BM) sites and the spleen of a mouse transplanted with a lentivirally barcoded human cord blood CD34+ HSPC xenograft. We observe high correlation between T cell samples across all anatomical sites, while B cell and Gr samples show high correlation to one another only within each anatomical site (**Fig. 2C**). This pattern is evident in the PCoA plot (**Fig. 2D**) as well. Unsorted cell samples from the spleen and pelvic BM vary in their clonal relationships to other samples, likely because of the underlying heterogeneity of the lineage composition of these bulk samples (**Fig. 2C**). These analyses support the notion that geographically isolated HSPC pools are responsible for the clonal composition of their respective geographic niches, and that the clonal composition of T cells across sites is largely independent from the output of these pools, supporting the thymic-dependent developmental pathway for T cells. Indeed, geographic isolation of HSPC output has been observed in another study in macaques; however, within the macaque study, T cell output early after transplantation was instead found to be anatomically compartmentalized and dependent at least short-term on these geographically isolated HSPC pools(25).

Comparing clonal distribution across animals from serial transplant experiments can also provide insight into the self-renewal capacity of engrafted, clonally marked cells. The Elder dataset contains clonal abundance information from serial xenograft mouse transplants of lentiviral-transduced ALL blasts. We observed high correlation of clonal abundances between samples collected from primary, secondary, and tertiary transplant recipient mice, excluding a few sites in the primary transplant (**Fig. 2E**). This aligns with the results presented by Elder et al(23) noting equipotential functional capability of ALL cells with some variation between sites, based on random sampling of the population of engrafted ALL cells. Samples from the same “generation” of serial transplant cluster together in principal coordinate space, supporting the notion presented in the study that ALL founder cells retain self-renewal capacity over several serial transplants (**Fig. 2F**). The distinct groupings of clones based on anatomic sampling site within the primary transplanted animals suggests that ALL clonal output also appears to be geographically compartmentalized, at least initially. Altogether, these three examples illustrate the utility of *barcodetrackR* in probing global clonal relationships between samples collected from various lineages or time points, providing valuable insights from a number of diverse biologic contexts.

### Determining clonality based on clonal counts and diversity measures

In clonal tracking studies, both clonal counts and diversity measures can provide insight into the clonality of progenitor cell pools. Here, we utilize *barcodetrackR* to assess clonality by visualizing the detected clone counts and Shannon diversities of samples from three datasets over time (**Fig. 3**). When quantifying clone numbers within the Six dataset, we show hundreds of unique integration sites retrieved across five purified peripheral blood lineage samples, with a larger number of clones detected in B cells and T cells as compared to Gr, Mo, and NK cells at most individual timepoints (**Fig. 3A**). Decreasing clonal diversity (Shannon index) in the NK cell lineage, as compared to other lineages, indicates that over time, a smaller number of clones account for a larger fraction of hematopoiesis in the NK cell compartment (**Fig 3B**). This implies a more oligoclonal population of contributing progenitors. The finding that the mature NK cell compartment is largely composed of a few high-contributing clones post-transplantation is in agreement with rhesus macaque barcoded autologous HSPC transplant studies(10).

**Figure 3:**
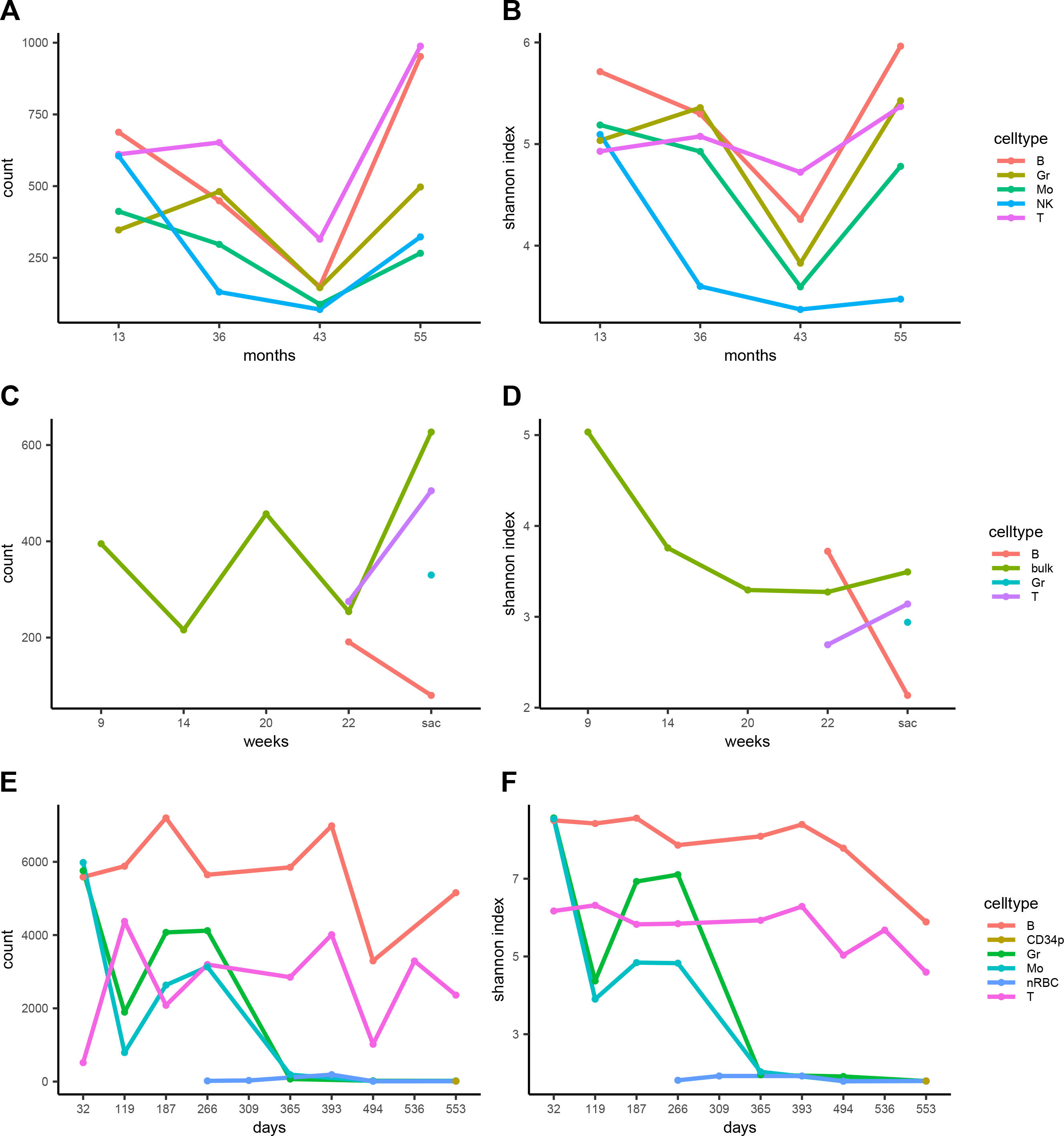
Measures of clonal diversity. Number of detected clones and Shannon diversity of clones in the Six, Belderbos, and Espinoza datasets. The number of clones detected in each lineage (**A**) and the Shannon diversity index of each sample (**B**) from the Six dataset. The number of clones detected in each lineage (**C**) and the Shannon diversity index of each sample (**D**) from the Belderbos dataset. The number of clones detected in each lineage (**E**) and the Shannon diversity index of each sample (**F**) from the Espinoza dataset. X-axes show months, weeks or days post-transplant with “sac” corresponding to the timepoint of euthanasia in the Belderbos dataset. The clone count reflects the number of unique clones detected in each sample, not the cumulative count at each timepoint. Shannon diversity is calculated on a per-sample basis based on the clonal population of each sample, not the cumulative number of clones. *Gr, Granulocyte; Mo, Monocyte; bulk, unsorted population; CD34p, CD34 positive cell; nRBC, nucleated Red Blood Cell*

Next, we analyzed patterns in the number of detected clones and Shannon diversities over time in peripheral blood samples from a single mouse xenograft obtained from the Belderbos dataset. We show that more clones were detected from the bulk peripheral blood sample at sacrifice than at the first time point (green line, **Fig. 3C**). The Shannon diversity of bulk samples decreased after the 9-week time point before stabilizing (**Fig. 3D**), underscoring the notion that clone counts alone are not ideal measures of sample diversity. These findings suggest that within this xenograft transplantation model, the diversity of HSPC output becomes stable over time, in agreement with previous long term clonal tracking studies in macaque(8) and human(7).

Finally, we use *barcodetrackR* to quantify an extreme case of minimal clonal counts and Shannon diversities in the context of clonal hematopoiesis using the Espinoza dataset (Table 1). In this study, multiple lentiviral insertions in an HSPC mediated dysplastic clonal erythroid and myeloid expansion, while largely sparing the lymphoid lineages. In agreement with the findings of the study, we find that the longitudinal clone numbers contributing to the B and T cell lineages fluctuate, but that the Shannon diversity index of these lineages remains high, especially at early time points, indicating polyclonal contribution to the lymphoid lineages (**Fig 3E**). However, after day 266 post-transplant, we observe a massive drop-off in both the number of unique clones detected (**Fig 3E**) and the Shannon diversity (**Fig 3F**) within the myeloid lineages. This coincides chronologically with the development of clonal hematopoiesis in the myeloid lineage. In the Espinoza study, 6 detected genetic tags in this dataset were all in fact found to correspond to a single lentivirally transduced cell with 9 insertions (3 of which were virtually undetected with the available dataset’s methodology due to deletions in the insertion proviral sequences), verifying that this pattern is representative of clonal hematopoiesis(21). While in the above examples we utilize unique detected clones at each time point, cumulative clone counts can also be calculated (**Fig. S1**) and provide a complementary view of clone numbers over time. Altogether, these examples emphasize the utility of diversity measures in interrogating the clonal output of progenitor pools in a number of contexts.

### Analyzing longitudinal clonal dynamics based on clonal abundances

Longitudinally tracking the abundance of individual clones can provide insight into clonal dynamics within lineages. We employed *barcodetrackR* to analyze longitudinal NK cell samples from an animal in the Wu et al study (Table 1), in which NK cell clonal dynamics were interrogated over 3 years in rhesus macaques receiving lentivirally barcoded autologous HSPCs(10). The detected clones in the NK cell compartment within this study remained largely independent from the HSPC pool responsible for the majority of non-NK hematopoiesis. We first visualize all individual NK cell clones from the Wu dataset in a binary heat map that depicts the presence or absence of all clones observed at 0.01% abundance or greater in at least one NK cell sample (**Fig 4A**). We find that new NK cell clones are detected at each time point, but that the number of newly detected clones decreases at later time points.

**Figure 4:**
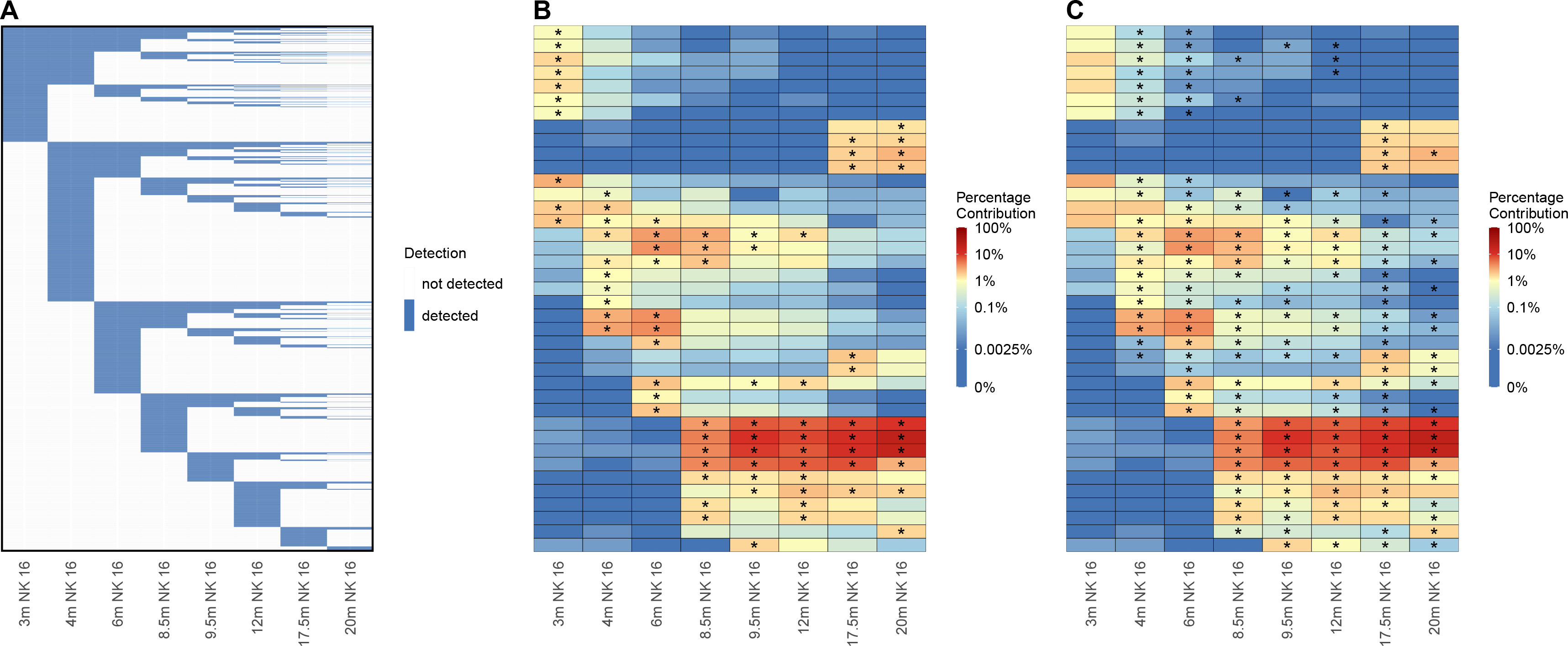
Longitudinal clonal patterns. Clonal dynamics of NK cells from the Wu dataset. (**A**) Binary heat map showing the presence (blue) or absence (white) of 11,799 individual clones detected at a proportion of 0.01% or greater in at least one NK cell sample from the Wu et al dataset. Columns represent samples and rows represent individual clones ordered by their first time point of detection. (**B**) Heat map showing the log normalized counts of the top ten clones from each NK cell sample from the Wu dataset. Overlaid asterisks indicate which clone is one of the top ten contributing clones for each sample, and clones are ordered on the y-axis based on hierarchical clustering of their Euclidean distances between their log normalized counts across samples. (**C**) Heat map depicting the same log normalized count values as in (**B**) but with overlaid asterisks instead indicating which clones significantly changed in proportion from the previous sample based on a p-value of < 0.05 assessed by a chi-squared test of proportions. *m, months post-transplant; NK 16, CD3-CD14-CD20-CD56-CD16+ NK cells*.

Next, we analyzed distinct clonal dynamics in individual NK cell clones using *barcodetrackR* to generate a heat map showing the abundance of the top ten NK cell clones from each sample over time (**Fig. 4B**). We visualize only the top clones in order to focus on the clones responsible for the majority of this cellular compartment’s clonal composition, with stars on the heatmap indicating the top ten contributing clones in each sample. One group of clones contributed at high levels for 3 months but subsequently declined. Of the top ten clones from the second time point (4 months), some clones declined in abundance while others increased in abundance at the 6-month time point before subsequently declining. A group clones became high abundance at 8.5 months and continued to contribute a large fraction of NK cell activity through the 20-month time point. Finally, a set of clones became high contributors at the 17.5-month time point after being virtually undetectable at previous timepoints. This analysis reveals the waxing and waning patterns of high-abundance NK cell clones over time, which were further interrogated in the Wu study. Lastly, we utilized statistical testing within *barcodetrackR* to view changes in proportion within the NK cell samples, marking clones with a star which had a statistically significant change (Chi-squared test) in abundance in the labelled sample in comparison to the previous sample (**Fig 4C**). This type of visualization and analysis further highlights the highly dynamic clonal patterns within the NK cell compartment. Altogether, these results indicate that the longitudinal tracking of highly abundant clones within datasets can provide insight into clonal dynamics at a single-clone level.

### Quantifying lineage bias based on shared clonality

Clonal tracking studies measure HSPC clonal contributions to different mature blood cell lineages. Thus, the lineage bias of HSPCs, such as those that skew towards myeloid or lymphoid lineages, can be interrogated on a global scale by comparing the ratio of clonal abundances between two specific lineages. Here, we use *barcodetrackR* to probe this concept of lineage bias in the Six clinical gene therapy trial dataset(6) and the Koelle rhesus macaque dataset(8)(Table 1), both based on lineage-purified samples following autologous transplantation with genetically tagged HSPCs.

We first track the density of clones, weighted by their added proportions, at each value of lineage bias (log ratio) between the Gr and T cell lineages across multiple timepoints in the Six dataset. We find the presence of three high-contributing sets of clones as determined by Gr/T lineage bias: Gr-biased (rightmost peak), balanced clones (middle peak), and T-biased (leftmost peak) (**Fig. 5A**). By systematically comparing each cell type in the dataset, we find that three sets of clones can be found when comparing Mo/T, Gr/B, or Mo/B lineages (**Fig. S2**) further supporting differences in upstream progenitors accounting for myeloid versus lymphoid lineages. In contrast, when comparing Gr/Mo or T/B lineages, we find that the majority of clones have balanced contribution to the two lineages (**Fig. S2**). Interestingly, clones contributing to the NK cell samples are predominantly unilineage, sharing very little clonality with other lineages, including other lymphoid lineages such as T and B cells (**Fig. S2**). This is in line with clonal tracking studies performed in a rhesus macaque animal model(2,10). Conducting the same analysis on longitudinal samples from the Koelle dataset reveals the presence of Gr-biased, balanced, and T-biased clones at the 4.5-month timepoint (**Fig. 5C**), consistent with the Six dataset. However, there is an increase in abundance of balanced clones at later timepoints, indicating a shift towards hematopoiesis from multipotent upstream progenitors, capable of reconstituting both myeloid and lymphoid lineages. This is also the case when comparing the T cell lineage to the Mo lineage as the majority of clones contribute similar abundances to Gr and Mo lineages (**Fig. S3**).

**Figure 5:**
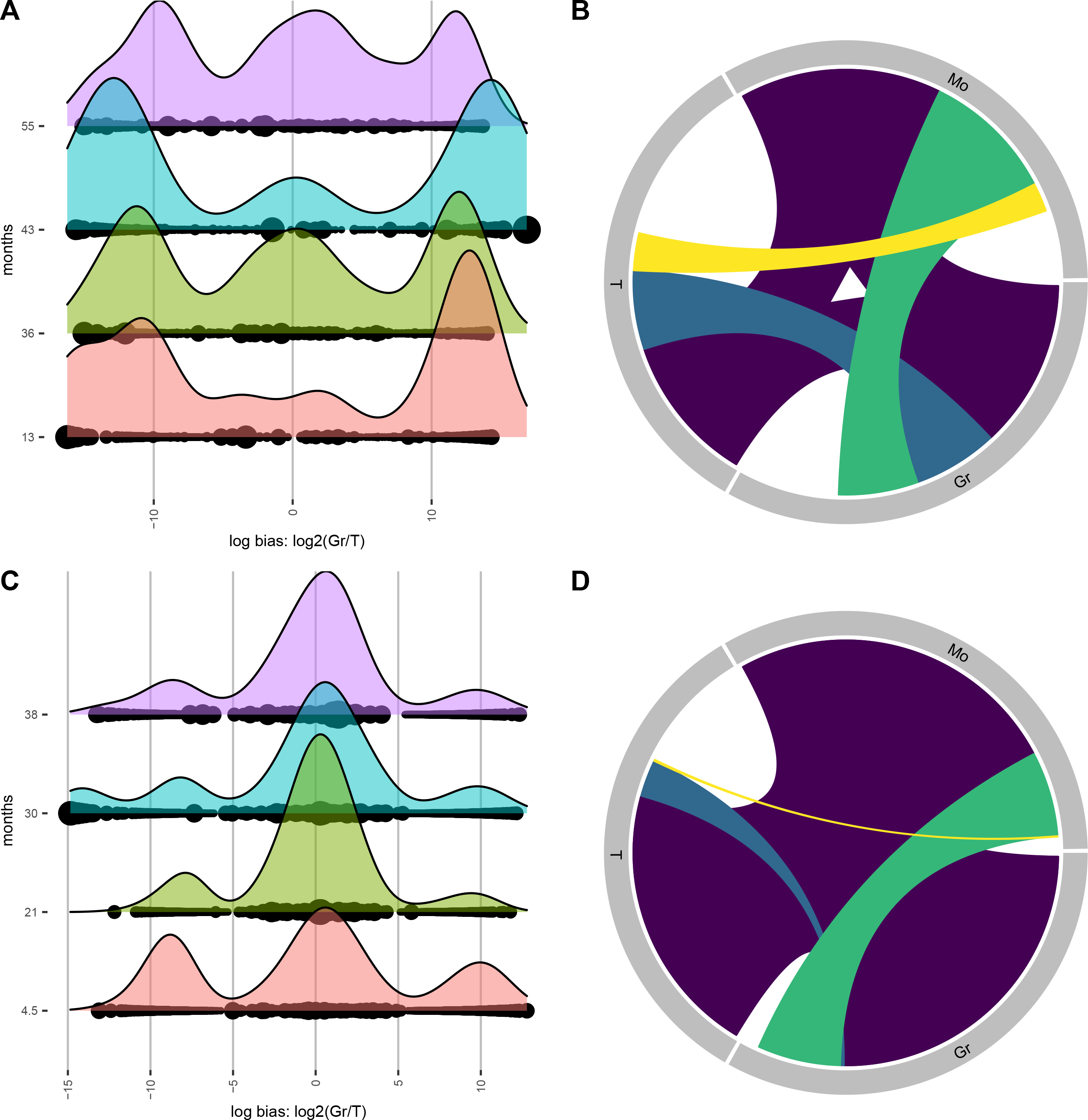
Lineage bias. Comparison of myeloid-lymphoid lineage bias in the Six and Koelle datasets. Ridge plot shows clonal bias between Gr and T lineages at multiple timepoints of the Six dataset (**A**) and the Koelle dataset (**C**). Ridges indicate the density of clones at the value of log-bias on the x-axes, and dots indicate individual clones, sized by their overall abundance. Multiple ridge plots along the y-axes correspond to the time point of each sample in months post-transplant. Chord diagram showing shared clonality between Gr, T, and Mo lineages from Six et al (**B**) and Koelle et al (**D**) datasets. Each uniquely colored chord represents a unique combination of lineages, and the width of each chord as it intersects with a lineage indicates the proportional contribution of that group of clones to that lineage. The space around the perimeter without a chord indicates the percentage contribution of unilineage clones. *Gr, Granulocyte; Mo, Monocyte*.

We next use *barcodetrackR* to construct an abundance-weighted chord diagram between three lineages in the Six dataset. We selected the Gr, T, and Mo lineages at the final 55-month time point (**Fig. 5B**), finding that a large fraction of detected hematopoiesis at this time point is shared between all three lineages (purple). However, there also exist clones detected only in two lineages (yellow, blue, green), and biased clones only found in one lineage, indicated by the white space around the perimeter. Likewise, a similar pattern is observed in the Koelle dataset at the final 38-month time point (**Fig. 5D**) with a large fraction of detected hematopoiesis arising from clones detected in all three lineages (purple). The fraction of detected hematopoiesis arising from T-Gr or T-Mo restricted clones (blue, yellow respectively), however, is minimal compared to that arising from Gr-Mo restricted clones (green), supporting the notion of upstream progenitors biased towards the myeloid lineage within this dataset. Thus, *barcodetrackR* provides a number of functions useful for inferring the lineage biases of upstream progenitors from clonal tracking data.

### barcodetrackR is versatile and includes a user-friendly graphical user interface

We highlight the versatility of *barcodetrackR* by analyzing clonal tracking data collected from TCR sequencing by visualizing the number of T cell clones detected and the Shannon diversity of longitudinal samples from X-SCID patients treated with HSPC gene therapy(24) (**Fig. S4**). The *barcodetrackR* package includes a graphical user interface (built using *shiny*(26)) to allow researchers without programming experience to use these advanced quantitative tools. After uploading genetic tag count matrices and metadata, users can toggle between tabs corresponding to different visualizations and analyses. Within each tab, users can specify calculation and display methods to create reproducible analyses and publication-quality data visualizations.

## DISCUSSION

A recent gathering of over 30 researchers in the clonal tracking field (2018 *StemCellMathLab* workshop(20)) formalized a call for the development of open-source tools for the analysis of clonal tracking data in order to promote rigor and reproducibility within the field. Here, we provide and showcase our open-source R package *barcodetrackR*, which encompasses an extensive, flexible, and accessible set of tools in order to address these needs and serve as a critical foundation on which to build further analytical approaches in the clonal tracking field. While tools for the processing of the raw sequencing data from clonal tracking experiments have been previously developed (16), *barcodetrackR* represents the first formal tool dedicated to interrogating the underlying biology represented by these clonal abundances. As shown, *barcodetrackR* is a multifaceted toolkit and has diverse applications, underscoring the utility of using complementary data analysis methods and visualizations to probe biological hypotheses. The development and implementation of a *shiny* app further adds to the utility of the package by making it more accessible to the clonal tracking community, which continues to expand as sequencing costs decline and methodologies continue to improve.

Although our package encompasses a large number of tools and methods, it is by no means an exhaustive toolbox, and we envision continuing to add to it in the future in order to address new biologic questions that arise. While the majority of prior clonal tracking experimental designs have precluded the acquisition of replicate samples and often encompassed few transplanted humans and/or animals, future studies will likely be able to acquire biological replicates in a number of different contexts to allow for more rigorous statistical testing of sample relationships and clonal dynamics. And while clustering methods to identify populations of clones with similar properties have thus far been limited to hierarchical(10) and k-means(6) in the literature, and to hierarchical clustering within *barcodetrackR*, the growing development of clustering frameworks of single cells in the scRNA-seq field may provide a future basis by which to identify clusters of clones based on longitudinal behavior and distribution across compartments(27). Furthermore, other challenges remain in the clonal tracking field, namely, the necessity for improved sharing and aggregating of data. We believe *barcodetrackR* can be of high utility to the clonal tracking field and serve as an important step towards building a more robust and reproducible analytical framework in the field.

## MATERIALS AND METHODS

### Package availability

The *barcodetrackR* package is freely available from GitHub under a Creative Commons 0 license and can be found at https://github.com/dunbarlabNIH/barcodetrackR. A frozen version of the package at the time of publication will be placed on Zenodo. The analytical and visualization tools included in *barcodetrackR* rely on the *ggplot2*(28), *vegan*(29), *proxy*(30), and *circlize*(31) packages, as well as a number of packages in the *tidyverse*(28) suite. The user interface is built using the *shiny* package in R(26). *barcodetrackR* can be installed in R using the *devtools*(28) utility and the command devtools::install_github(“dunbarlabNIH/barcodetrackR”). Within the GitHub repository, we include a vignette illustrating the use of *barcodetrackR* on several published barcoding datasets, which is available as an R markdown file or in html format online at: https://dunbarlabNIH.github.io/barcodetrackR. All figures included in the manuscript were generated using the *barcodetrackR* package, and the R markdown file “create_all_figures.Rmd” is included in the *barcodetrackR* GitHub.

### Dataset availability

The datasets used in this study are publicly available from published barcoding experiments and detailed in Table 1. Data were pre-processed in R to create tabular data files amenable for entry into *barcodetrackR*, and these procedures are outlined in scripts within the inst/sample_data directory within the *barcodetrackR* package on GitHub.

### Data collection, genetic tag retrieval, and normalization

Multiple clonal tracking methodologies exist in the literature(30), with the most recent methods relying on next-generation sequencing to retrieve lineage tracing elements. Several analysis pipelines exist for the retrieval and error-correction of lineage tracing elements from sequencing data(14–17). The experimental techniques utilized, the number of cells sampled, the level of tagged cell within the population, the sequencing platform applied, and the computational method of genetic tag extraction affect the number and frequency of tags detected in a lineage tracing study.

The *barcodetrackR* package can operate on any dataset that contains rows as observations and columns as samples, regardless of which experimental method for genetic labelling and approaches for tag retrieval were used. When creating a *SummarizedExperiment* (SE) object from the publicly available *Bioconductor* repository(32) for input into *barcodetrackR*, users have the option of including a threshold to exclude low-abundance occurrences that are more likely to come from noise or sequencing error(2). Using a threshold of 0.005, for example, retains genetic tags which are present at an abundance of 0.5% or greater in at least one sample. Within this paper, the Six, Elder, Espinoza, Wu, and Koelle datasets were used with a threshold of 0.0001. The Belderbos and Clarke datasets were used with no threshold.

Instantiating a SE through the function *create_SE* automatically creates the following assays: *counts*: the raw values from the input dataframe, *percentages*: the per-column proportions of each entry in each column, *ranks*: the rank of each entry in each column, *normalized*: the normalized read values in counts-per million (CPM), and *logs*: the log of the normalized values. The default normalization is counts per million, and log-normalized values are calculated by taking the log of plus-one normalized data so that zeros are retained. Both the scale factor and the log base can be passed as arguments to the *create_SE* command. The use of an SE object permits the addition of custom assays to the object to facilitate flexibility (e.g. custom normalization strategies).

### Global clonal distributions

Users can view correlations between samples on a grid using the *cor_plot* function, longitudinally using the *autocor_plot* function, or for two samples using the *scatter_plot* function. Choices for correlation measures include “*pearson*”, “*spearman*”, or “*kendall*.” The functions include display parameters as inputs, such as whether to display correlation values in a *cor_plot* grid as color, colored circles, or showing the actual correlation value.

The *mds_plot* function calculates dissimilarity indices between samples using any distance metric within the *vegdist* function from the R package *vegan*(29). At time of publication, these include “*manhattan*”, “*euclidean*”, “*canberra*”, “*clark*”, “*bray*”, “*kulczynski*”, “*jaccard*”, “*gower*”, “*altGower*”, “*morisita*”, “*horn*”, “*mountford*”, “*raup*”, “*binomial*”, “*chao*”, or “*cao*”. The distance methods produce a matrix composed of the distances between samples, given the composition of genetic tags in each sample. Principal coordinates analysis is performed on the distance matrix using the base R function *cmdscale* in order to display dissimilarity between samples in two dimensions.

### Clonal diversity

Three measures of within-sample diversity can be calculated by the function *clonal_diversity*: shannon diversity (H’), simpson diversity (λ), and inverse-simpson, which is calculated as 1/λ. Their equations are as follows:

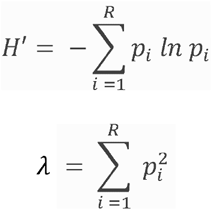

Where *R* is the total number of species (in this case genetic tags), and *p_i_* is the proportion of each gentic tag in the sample. Additionally, users can simply display the nominal count or Shannon count. Shannon count is calculated as:

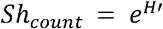

To accurately compare diversity between samples, the same number of labeled cells should be used as starting material for quantification. This is especially important when assessing diversity based on the nominal count of genetic tags retrieved. Belderbos et al showed that the Shannon count is stable with respect to filtering thresholds and stays below the theoretical library size upon re-sampling(22). Therefore, in some cases, it may be beneficial to use Shannon count rather than nominal genetic tag counts when comparing diversity between samples.

### Clonal patterns

Heat maps created by the function *barcode_ggheatmap* display clonal abundances across samples by coloring cells based on the log-normalized abundance of each clone. Users specify the ordering of samples along the x-axis and the number of clones displayed per sample. By default, the top clones from each sample are marked by a star, but the user can also choose to label each entry in the heatmap by percentage or number of reads. The function includes aesthetic parameters, such as the label size, percent scale, and color scale, which can also be easily controlled within the *barcodetrackR* graphical user interface.

Under default settings for the heat map function, individual clones are hierarchically clustered along the y-axis based on their log-abundance (or other specified assay) across samples. Setting the “*dendro*” parameter to TRUE allows users to see this clustering and setting the “*clusters*” parameter to a non-zero value will label hierarchical clusters of clones. The dendrogram is drawn using *ggdendro*(33). Users can choose from a number of methods to calculate the distance metric (any method included in the proxy R package) and clustering (“*ward.D*”, “*ward.D2*”, “*single*”, “*complete*”, “*average*”, “*mcquitty*”, “*median*”, or “*centroid*”). Clones can also be ordered by the first sample in which they had a non-zero abundance by setting the “*row_order*” parameter to “*emergence*.”

To track the emergence of clones over time, the function *barcode_binary_heatmap* will display a heatmap indicating the presence or absence of clones in each sample. The “*threshold*” parameter to the function dictates the limit of detection. Clones with percentage abundance below this threshold in a given sample will be set to “absent.” Clones are ordered in the binary heatmap based on the first sample in which they emerged (had a non-zero abundance).

The function *barcode_ggheatmap_stat* allows users to quantify changes in clone abundance based on statistical testing. This function requires the sample size of cells for each sample which cannot be calculated from the genetic tag count data. The sample size should be the number of labeled cells before amplification, because this is the population of cells which the clonal tracking data represent. To compare barcode abundances between samples, users can choose from a “*chi-squared*” or “*fisher*” exact test. The tests operate on a contingency table for each clone to determine whether the clone changed in proportion based on the abundance and sample size. By default, each sample is compared to the previous, but users can also specify to compare each sample to a single reference sample (such as the initial time point) by setting the “*stat_option*” parameter to “*reference*.” The user can specify a p-value threshold to assign significance and can choose to display p-values on the heat map, rather than stars indicating statistically significant changes. Also, the user can specify to only show clones which increase or decrease in proportion through the “*stat_display*” parameter.

### Clonal bias

The *ridge_plot* function calculates bias as a continuous variable and displays the density of clones at each level of log bias. In order to calculate log bias between clones that are only present in one sample, log bias is calculated as followed within the ridge plot function:

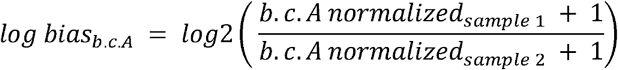

The normalized value is taken from the SE slot which is scaled to counts per million. The option to only analyze clones present in both samples is also included in the *ridge_plot* function through the parameter “*remove_unique*”. The density of clones at each value of log bias is estimated using the kernel density estimator included in the *ggplot2* R package. When the “*weighted*” parameter is set to TRUE, the function weights the density estimation by the *combined abundance* of the clone between the two samples, calculated as:

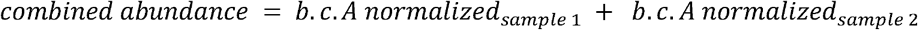

### Chord diagrams

The *circos_plot* function utilizes the circlize package in R(31) to display shared clones between samples. Samples are shown as regions around a circle with their shared clonality shown as links between regions. With each unique combination, a new link is created with a unique color drawn from a sequential color palette. When the “*weighted*” option is set to “*FALSE*”, the function operates on the *counts* assay. Therefore, the length of each region around the circle represents the number of clones detected in each sample, and the width of links between regions is proportional to the number of shared clones. When the “*weighted*” option is set to “*TRUE*”, the function operates on the *percentages* slot. Each region around the circle has the same length corresponding to 100%, and the links between regions correspond to the fractional abundance of the shared clones within each sample. Therefore, when the “*weighted*” option is set to “*TRUE*”, the same link can have a different width at each connection to a region.

## DECLARATIONS

### Ethics approval and consent to participate

Not applicable.

### Consent for publication

Not applicable.

### Availability of data and materials

Accessibility to all datasets used is outlined in Table 1. *barcodetrackR* is an open-source R package on GitHub at https://github.com/dunbarlabNIH/barcodetrackR and the current version of the R package will be frozen on Zenodo at the time of publication.

### Authors’ contributions

DAE and RDM wrote the manuscript. DAE and RDM developed code and performed analysis of existing datasets. SJK and CW aided with development of visualizations. CED supervised the project and edited the manuscript.

## Acknowledgements

The authors would like to thank David Allan of the NHLBI Intramural Research Program for his contributions to approaches for statistical testing of barcode abundances. The authors thank members of the Dunbar lab for helpful feedback in revision of this manuscript.

---

**Figure S1:**
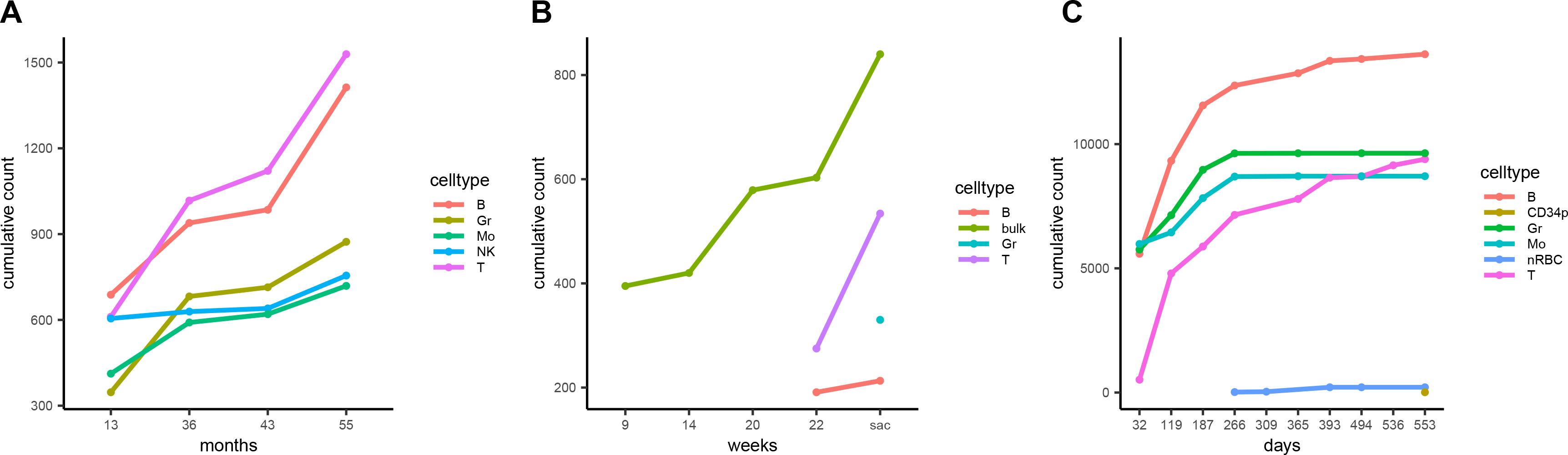
Cumulative clone counts from the Six, Belderbos, and Espinoza datasets. The cumulative count of unique clones detected in each lineage across longitudinal samples from the Six (**A**), Belderbos (**B**), and Espinoza (**C**) datasets. X-axes show months, weeks or days post-transplant with “sac” corresponding to the timepoint of euthanasia in the Belderbos dataset. *Gr, Granulocyte; Mo, Monocyte; bulk, unsorted population; CD34p, CD34 positive cell; nRBC, nucleated Red Blood Cell*

**Figure S2:**
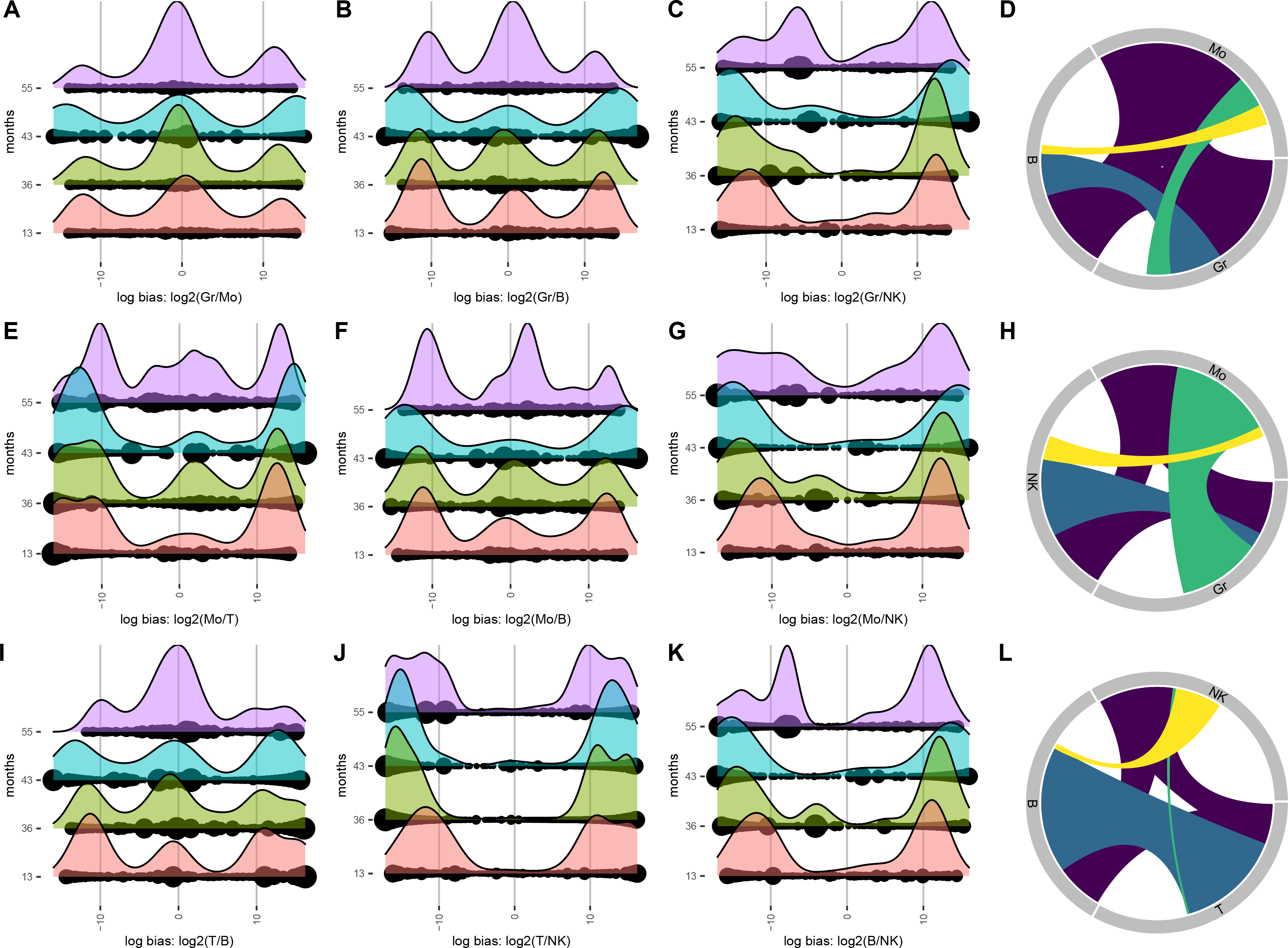
Systematic analysis of lineage bias in the Six dataset. Ridge plots comparing the lineage bias at multiple time points between all pairwise combinations of lineages from the Six dataset not shown in Fig. 5 (**A-C, E-G, I-K**). Ridges indicate the density of clones at the value of log-bias on the x-axes, and dots indicate individual clones, sized by their overall abundance. Multiple ridge plots along the y-axes correspond to the time point of each sample in months post-transplant. Chord diagram showing shared clonality between Gr, B, and Mo lineages (**D**), Gr, NK, and Mo lineages (**H**), and T, B, and NK lineages (**L**) from Six et al. Each uniquely colored chord represents a unique combination of lineages, and the width of each chord as it intersects with a lineage indicates the proportional contribution of that group of clones to that lineage. The space around the perimeter without a chord indicates the percentage contribution of unilineage clones. *Gr, Granulocyte; Mo, Monocyte; NK, Natural Killer cell*.

**Figure S3:**
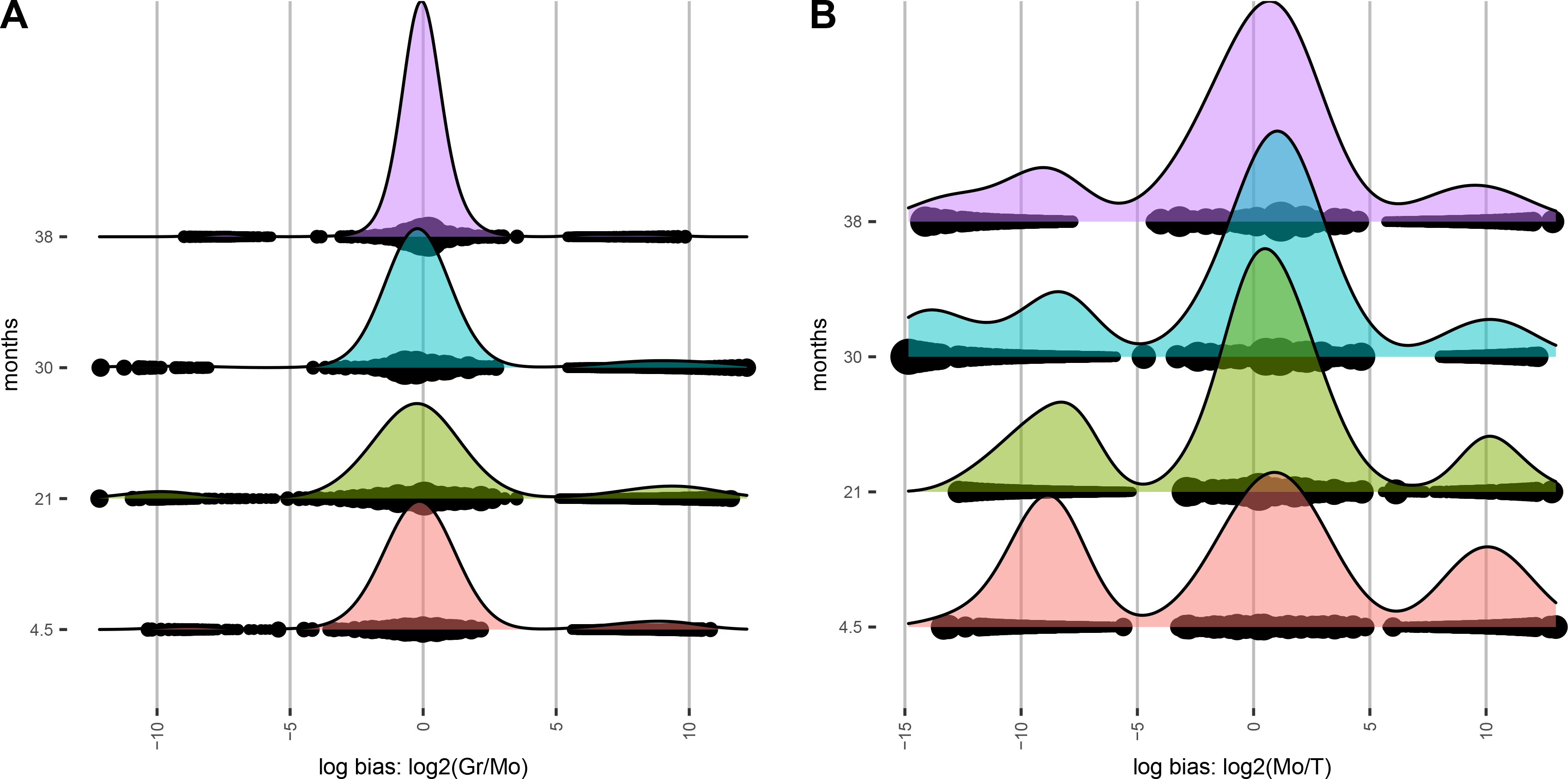
Systematic analysis of lineage bias in the Koelle dataset. Ridge plots comparing the lineage bias at multiple time points between Gr and Mo lineages (**A**) and Mo and T lineages (**B**) from the Koelle dataset. Comparison of the Gr and T lineage is shown in Fig. 5. Ridges indicate the density of clones at the value of log-bias on the x-axes, and dots indicate individual clones, sized by their overall abundance. Multiple ridge plots along the y-axes correspond to the time point of each sample in months post-transplant. *Gr, Granulocyte; Mo, Monocyte*.

**Figure S4:**
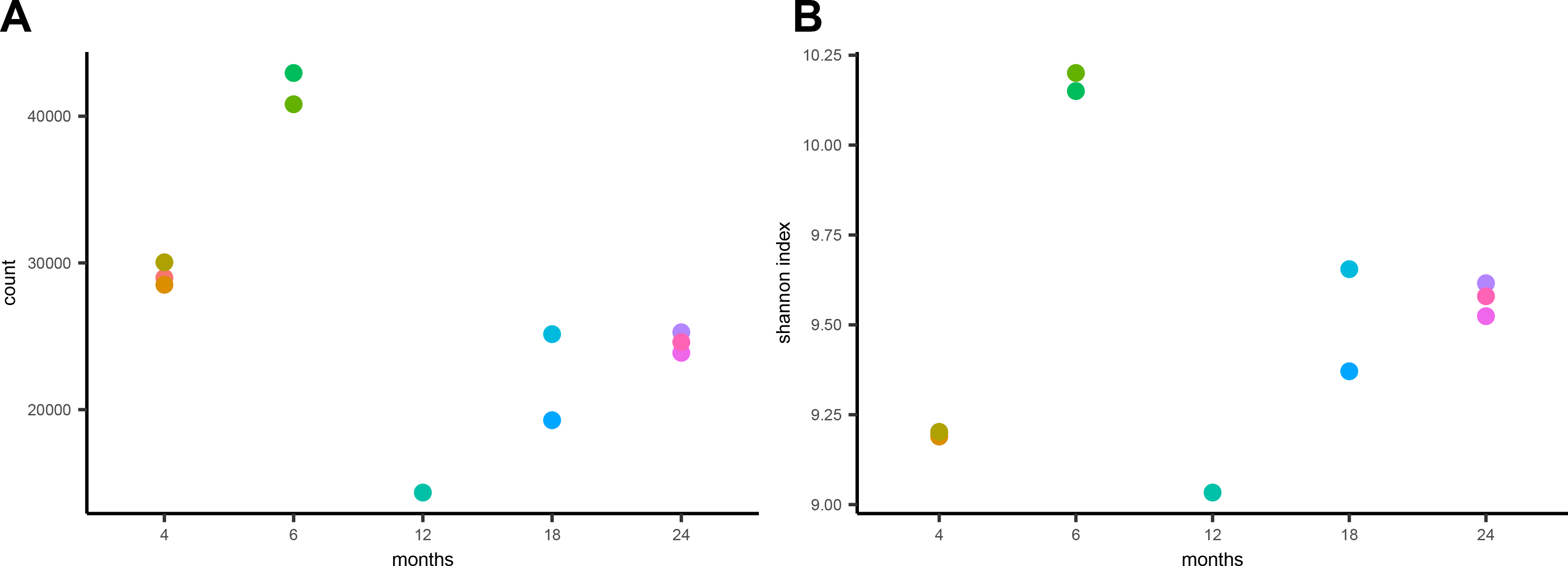
Clonal diversity from TCR sequencing data. The number of T cell clones detected at each timepoint (**A**) and the Shannon diversity (**B**) of T cell clones in each sample from the Clarke et al dataset (Table 1). Clone counts represent the number of unique TCR sequences detected at each point not the cumulative number of clones. Multiple replicates are shown at the 4, 6, 18, and 24 month time points, as described in Clarke et al(24). Each color represents an independent replicate. The x-axes represent months post-transplant.

## REFERENCES

1. Lu R, Neff NF, Quake SR, Weissman IL. Tracking single hematopoietic stem cells in vivo using high-throughput sequencing in conjunction with viral genetic barcoding. Nat Biotechnol. 2011 Oct;29(10):928–33.

2. Wu C, Li B, Lu R, Koelle SJ, Yang Y, Jares A, et al. Clonal Tracking of Rhesus Macaque Hematopoiesis Highlights a Distinct Lineage Origin for Natural Killer Cells. Cell Stem Cell. 2014 Apr;14(4):486–99.

3. Radtke S, Adair JE, Giese MA, Chan Y-Y, Norgaard ZK, Enstrom M, et al. A distinct hematopoietic stem cell population for rapid multilineage engraftment in nonhuman primates. Sci Transl Med. 2017 Nov 1;9(414):eaan1145.

4. Kim S, Kim N, Presson AP, Metzger ME, Bonifacino AC, Sehl M, et al. Dynamics of HSPC Repopulation in Nonhuman Primates Revealed by a Decade-Long Clonal-Tracking Study. Cell Stem Cell. 2014 Apr;14(4):473–85.

5. Gerrits A, Dykstra B, Kalmykowa OJ, Klauke K, Verovskaya E, Broekhuis MJC, et al. Cellular barcoding tool for clonal analysis in the hematopoietic system. Blood. 2010 Apr 1;115(13):2610–8.

6. Six E, Guilloux A, Denis A, Lecoules A, Magnani A, Vilette R, et al. Clonal tracking in gene therapy patients reveals a diversity of human hematopoietic differentiation programs. Blood. 2020 Apr 9;135(15):1219–31.

7. Biasco L, Pellin D, Scala S, Dionisio F, Basso-Ricci L, Leonardelli L, et al. In Vivo Tracking of Human Hematopoiesis Reveals Patterns of Clonal Dynamics during Early and Steady-State Reconstitution Phases. Cell Stem Cell. 2016 Jul;19(1):107–19.

8. Koelle SJ, Espinoza DA, Wu C, Xu J, Lu R, Li B, et al. Quantitative stability of hematopoietic stem and progenitor cell clonal output in rhesus macaques receiving transplants. Blood. 2017 Mar 16;129(11):1448–57.

9. Brugman MH, Wiekmeijer A-S, van Eggermond M, Wolvers-Tettero I, Langerak AW, de Haas EFE, et al. Development of a diverse human T-cell repertoire despite stringent restriction of hematopoietic clonality in the thymus. Proc Natl Acad Sci USA. 2015 Nov 3;112(44):E6020–7.

10. Wu C, Espinoza DA, Koelle SJ, Yang D, Truitt L, Schlums H, et al. Clonal expansion and compartmentalized maintenance of rhesus macaque NK cell subsets. Sci Immunol. 2018 Nov 2;3(29):eaat9781.

11. Merino D, Weber TS, Serrano A, Vaillant F, Liu K, Pal B, et al. Barcoding reveals complex clonal behavior in patient-derived xenografts of metastatic triple negative breast cancer. Nature Communications. 2019 Feb 15;10(1):1–12.

12. Porter SN, Baker LC, Mittelman D, Porteus MH. Lentiviral and targeted cellular barcoding reveals ongoing clonal dynamics of cell lines in vitro and in vivo. Genome Biology. 2014 May 30;15(5):R75.

13. Sheih A, Voillet V, Hanafi L-A, DeBerg HA, Yajima M, Hawkins R, et al. Clonal kinetics and single-cell transcriptional profiling of CAR-T cells in patients undergoing CD19 CART immunotherapy. Nature Communications. 2020 Jan 10;11(1):1–13.

14. Berry CC, Nobles C, Six E, Wu Y, Malani N, Sherman E, et al. INSPIIRED: Quantification and Visualization Tools for Analyzing Integration Site Distributions. Molecular Therapy - Methods & Clinical Development. 2017 Mar;4:17–26.

15. Sherman E, Nobles C, Berry CC, Six E, Wu Y, Dryga A, et al. INSPIIRED: A Pipeline for Quantitative Analysis of Sites of New DNA Integration in Cellular Genomes. Molecular Therapy - Methods & Clinical Development. 2017 Mar;4:39–49.

16. Thielecke L, Cornils K, Glauche I. genBaRcode: a comprehensive R-package for genetic barcode analysis. Wren J, editor. Bioinformatics. 2020 Apr 1;36(7):2189–94.

17. Bramlett C, Jiang D, Nogalska A, Eerdeng J, Contreras J, Lu R. Clonal tracking using embedded viral barcoding and high-throughput sequencing. Nat Protoc. 2020 Apr;15(4):1436–58.

18. Wolf FA, Angerer P, Theis FJ. SCANPY: large-scale single-cell gene expression data analysis. Genome Biol. 2018 Dec;19(1):15.

19. Stuart T, Butler A, Hoffman P, Hafemeister C, Papalexi E, Mauck WM, et al. Comprehensive Integration of Single-Cell Data. Cell. 2019 Jun;177(7):1888–1902.e21.

20. Lyne A-M, Kent DG, Laurenti E, Cornils K, Glauche I, Perié L. A track of the clones: new developments in cellular barcoding. Experimental Hematology. 2018 Dec;68:15–20.

21. Espinoza DA, Fan X, Yang D, Cordes SF, Truitt LL, Calvo KR, et al. Aberrant Clonal Hematopoiesis following Lentiviral Vector Transduction of HSPCs in a Rhesus Macaque. Molecular Therapy. 2019 Jun 5;27(6):1074–86.

22. Belderbos ME, Jacobs S, Koster TK, Ausema A, Weersing E, Zwart E, et al. Donor-to-Donor Heterogeneity in the Clonal Dynamics of Transplanted Human Cord Blood Stem Cells in Murine Xenografts. Biology of Blood and Marrow Transplantation. 2020 Jan 1;26(1):16–25.

23. Elder A, Bomken S, Wilson I, Blair HJ, Cockell S, Ponthan F, et al. Abundant and equipotent founder cells establish and maintain acute lymphoblastic leukaemia. Leukemia. 2017 Dec;31(12):2577–86.

24. Clarke EL, Connell AJ, Six E, Kadry NA, Abbas AA, Hwang Y, et al. T cell dynamics and response of the microbiota after gene therapy to treat X-linked severe combined immunodeficiency. Genome Med. 2018 Dec;10(1):70.

25. Wu C, Espinoza DA, Koelle SJ, Potter EL, Lu R, Li B, et al. Geographic clonal tracking in macaques provides insights into HSPC migration and differentiation. J Exp Med. 2018 Jan 2;215(1):217–32.

26. Chang W, Cheng J, Allaire JJ, Xie Y, McPherson J. shiny: Web Application Framework for R [Internet]. 2020. Available from: https://CRAN.R-project.org/package=shiny

27. Kiselev VY, Andrews TS, Hemberg M. Challenges in unsupervised clustering of single-cell RNA-seq data. Nat Rev Genet. 2019 May;20(5):273–82.

28. Wickham H, Averick M, Bryan J, Chang W, McGowan LD, François R, et al. Welcome to the Tidyverse. Journal of Open Source Software. 2019 Nov 21;4(43):1686.

29. Oksanen J, Blanchet FG, Friendly M, Kindt R, Legendre P, McGlinn D, et al. vegan: Community Ecology Package [Internet]. 2019. Available from: https://CRAN.R-project.org/package=vegan

30. Meyer D, Buchta C. proxy: Distance and Similarity Measures [Internet]. 2020. Available from: https://CRAN.R-project.org/package=proxy

31. Gu Z, Gu L, Eils R, Schlesner M, Brors B. circlize implements and enhances circular visualization in R. Bioinformatics. 2014 Oct 1;30(19):2811–2.

32. Morgan M, Obenchain V, Hester J, Pagès H. SummarizedExperiment: SummarizedExperiment container [Internet]. 2020. Available from: https://bioconductor.org/packages/SummarizedExperiment/

33. de Vries A, Ripley BD. ggdendro: Create Dendrograms and Tree Diagrams using “ggplot2” [Internet]. 2016. Available from: https://CRAN.R-project.org/package=ggdendro

